# The Neurochemical Mechanisms Underlying the Enhancing Effects of Rewards and Punishments on Motor Performance

**DOI:** 10.1101/2023.03.16.532906

**Authors:** R. Hamel, J. Pearson, L. Sifi, D. Patel, M.R. Hinder, N. Jenkinson, J.M. Galea

## Abstract

Monetary rewards and punishments enhance motor performance and are associated with corticospinal excitability (CSE) increases within the motor cortex (M1) during movement preparation. However, such CSE changes have unclear origins; they could stem from increased glutamatergic (GLUTergic) facilitation and/or decreased type A gamma-aminobutyric acid (GABA_A_)-mediated inhibition within M1. To investigate this, paired-pulse transcranial magnetic stimulation was used to assess GLUTergic facilitation and GABA_A_ inhibition within M1 whilst participants prepared to execute 4-element finger-press sequences. Behaviourally, rewards and punishments enhanced both reaction and movement times. Neurochemically, regardless of rewards or punishments, a digit-*specific* increase in GLUTergic facilitation and digit-*unspecific* decrease in GABA_A_ inhibition occurred during preparation as movement onset approached. In parallel, both rewards and punishments non-specifically increased GLUTergic facilitation, but only rewards non-specifically decreased GABA_A_ inhibition during preparation. This suggests that, to enhance performance, rewards both increase GLUTergic facilitation and decrease GABA_A_ inhibition whilst punishments selectively increase GLUTergic facilitation. A control experiment revealed that such changes were not observed post-movement as participants processed reward and punishment feedback, indicating they were selective to movement preparation. Collectively, these results map the neurochemical changes in M1 by which incentives enhance motor performance.

## Introduction

Monetary rewards and punishments are known to enhance motor performance^1,2^ by facilitating the processes underlying movement preparation^3–11^. For instance, Freeman and Aron (2014)^6^ used transcranial magnetic stimulation (TMS) over the primary motor cortex (M1) to show that rewarded stimuli, as compared to neutral ones, quickened reaction time (RT) and increased corticospinal excitability (CSE) during movement preparation. Whilst this indicates that brain activity changes during movement preparation are associated with incentive-induced (reward/punishment) improvements in motor performance, the origins of such CSE changes remain unclear. Namely, changes in CSE reflect the excitability of cortical, subcortical, as well as spinal structures^12^, which can further reflect an increase in glutamatergic (GLUTergic) and/or a decrease in gamma-aminobutyric acid (GABA)ergic activity^13,14^. Importantly, rewards and punishments are increasingly recognised as potential enhancers of rehabilitation following physical and brain insults^15–18^. Therefore, elucidating their mechanisms of action could lead to improved therapeutic interventions^19^. This work sought to characterise the circuit-specific changes occurring within M1 when motor performance is enhanced by rewards and punishments.

Converging lines of evidence suggest that rewards and punishments enhance performance by altering M1’s intracortical GLUTergic and type A GABA (GABA_A_) activity during movement preparation. First, human anatomical evidence shows that ∼ 67% of M1’s neurons are projecting GLUTergic neurons whilst ∼ 33% of M1’s neurons are local GABA_A_ interneurons^20^. Assuming that CSE changes induced by rewards and punishments have some cortical origins, one tentative possibility is that they originate from intracortical GLUTergic neurons and GABA_A_ interneurons in M1. To date, no study has directly evaluated the effects of rewards and punishments on GLUTergic and GABA_A_ circuits in M1 during movement preparation. Nonetheless, human TMS studies have shown that movement preparation releases M1’s intracortical GABA_A_-mediated inhibition^21–23^, pointing to the role of local GABA_A_ interneurons in mediating movement preparation. Whether movement preparation entails changes in M1’s GLUTergic activity remains unclear in humans but this proposition is supported by animal work^24,25^. Overall, this evidence suggests that any enhancement of performance induced by rewards and punishments must be reflected in M1’s GLUTergic and GABA_A_ activity during movement preparation. Second, parallel evidence has shown that the ventral tegmental area (VTA) alters M1’s excitability^26^ through GLUTergic^27^ and dopaminergic signalling^28^ by activating type 2 dopamine (D2)-like receptors^29^ located on GABA_A_ (parvalbumin-expressing) interneurons^30^. Given the well-established role of VTA neurons in processing both rewards and punishments^31,32^, this pathway could underpin changes in GLUTergic and GABA_A_-mediated inhibition in M1 that lead to the invigorating effects of incentives on motor performance. In further support of this, pharmacological work has shown that memantine^33^ (a N-methyl-d-Aspartate [NMDA] receptor antagonist), diazepam^34^ (a GABA_A_ agonist) and ethyl alcohol^35^ (a NMDA antagonist and GABA_A_ agonist) alter behavioural responses to rewards and punishments. Collectively, this evidence suggests that incentives enhance motor performance through circuit-specific changes in GLUTergic and GABA_A_-mediated inhibition within M1. The objective of this work was to test this hypothesis.

To this end, neuronavigated paired-pulse TMS (ppTMS) was used to evaluate intracortical facilitation (ICF) and short intracortical inhibition (SICI) within the left M1 as participants prepared to execute finger-press sequences with their right hand (Figure 1A). ICF and SICI are well-established measures of GLUTergic facilitation and GABA_A_-mediated inhibition within M1^14,36^, respectively, indicating that ppTMS can reliably assess changes in these intracortical circuits without any confounding influences of changes within subcortical or spinal structures^14,36^. The ppTMS pulses were delivered at three time points to assess ICF and SICI changes during a Movement Preparation session (Figure 1A). To determine if these changes were specific to movement preparation, ICF and SICI were also assessed following the delivery of rewards and punishments in a Feedback Processing session (Figure 3A). Whilst each participant took part in both the Movement Preparation and Feedback Processing sessions, ICF and SICI were evaluated in separate groups of 20 participants (Figures 1C and 1D; Figures 3C and 3D). To investigate motor performance, participants were cued to execute 4-element finger-press sequences as fast and accurately as possible during Reward (max: +£0.6; min: +£0.0), Neutral (max: +£0.0; min: +£0.0), and Punishment conditions (max: +£0.0; min: -£0.6). To determine if ICF and SICI changes were (task-) digit-*specific*, participants executed sequences comprising index and little finger presses only (i.e., no middle or ring finger; Figure 1A) whilst ppTMS targeted M1’s cortical representation of the first dorsal interosseus (FDI), a co-agonist of for index finger flexion required for button presses^37^. Observing ICF or SICI changes in the FDI electromyography (EMG) data specifically when the index finger initiated a sequence would imply (task-) digit-*specific* activity in M1. Conversely, observing ICF and SICI changes in FDI EMG data that manifest regardless of the index initiating the sequence would suggest (task-) digit-*unspecific* activity in M1.

**Figure 1.**
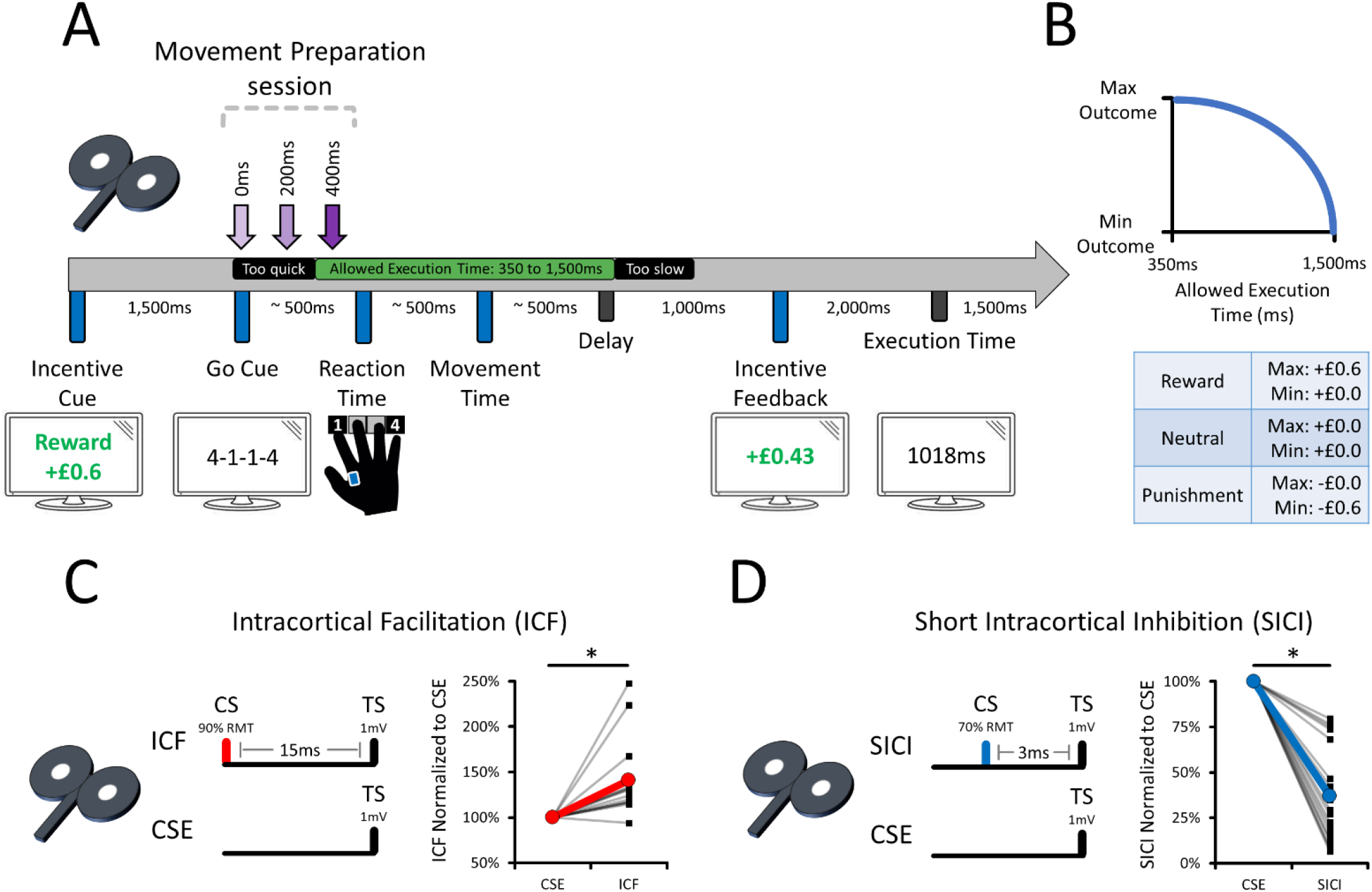
Assessment of both ICF and SICI during the Movement Preparation session. **(A)** *Chronology of a typical trial*. For both ICF and SICI, TMS was delivered either at GoCue onset (i.e., 0ms), 200ms or 400ms later. The TMS data were later pooled according to the percentage of the RT at which pulses were delivered (< 50% RT for the Early RT and > 50% RT for Late RT). Although participants executed the sequences by using their index and little fingers, TMS-evoked EMG responses were recorded from the FDI muscle only. **(B)** *The decay function used to adjust the incentives based on performance*. “Max” and “Min” refer to the maximal and minimal possible outcome on a given trial. Allowed execution time is defined as the sum of RT and movement time (MT; as depicted in panel A). **(C)** *The TMS parameters for the ICF Experiment (n = 20)* successfully induced ICF at rest, as assessed by delivering 30 single pulse and 30 paired ICF pulses. **(D)** *The TMS parameters for the SICI Experiment (n = 20)* successfully induced SICI at rest, as assessed by delivering 30 single pulses and 30 paired SICI pulses. For panels (C) and (D), individuals’ data with their respective means are shown^38^. RMT means resting motor threshold.

## RESULTS – MOVEMENT PREPARATION

### ICF and SICI were reliably observed at rest

To ensure the chosen TMS parameters reliably induced ICF and SICI (Figure 1C and 1D), 30 TMS trials were recorded for CSE, ICF, and SICI at rest before participants executed the finger-press sequences. The results revealed significant M1 facilitation with ICF (Figure 1C; 1.265 ± 0.155 mV) as compared to CSE (0.912 ± 0.111 mV; β = 0.3525 ± 0.0725, *p* < 0.0001). In addition, there was significant M1 inhibition with SICI (Figure 1D; 0.328 ± 0.074 mV) as compared to CSE (1.051 ± 0.161 mV; β = -0.7223 ± 0.1297, *p* < 0.0001). Note that Figure 1C and 1D reports ICF and SICI values normalised (%) to the average of CSE trials within each participant. Overall, this confirms that reliable ICF and SICI was observed at rest across participants.

### Digit-*specific* CSE increases during preparation

Generalised mixed linear models, using a gamma distribution to account for the positive continuous skewness in the EMG data^39^, were conducted to investigate CSE changes during movement preparation. Unless otherwise indicated, all generalised mixed linear models conducted in this work used a gamma distribution^39^. The fixed effects included in the models were Incentives (Reward, Neutral, Punishment), Time Points (GoCue, Early RT, Late RT), and Initiating Finger (Index, Little). Only the effects and interactions involving Time Points and Initiating Finger are reported in this section.

For the ICF experiment, the results revealed a Time Points * Initiating Finger interaction (χ^2^ = 20.890; *p* < 0.0001). Breakdown of the Time Points * Initiating Finger revealed a simple effect of Time Points for both the Index (χ^2^ = 8.824; *p* = 0.0121) and Little fingers (χ^2^ = 11.698; *p* = 0.0029). For the Index finger (not shown on Figure 2), the results revealed an *increase* in CSE at Late RT (2.203 ± 0.288 mV) as compared to Early RT (1.800 ± 0.274 mV; β = 0.4022 ± 0.1354; *p* = 0.0045) and GoCue (1.838 ± 0.290 mV; β = 0.3645 ± 0.1487; *p* = 0.0213). CSE measured at Early RT and GoCue did not significantly differ (β = 0.0377 ± 0.0873; *p* = 0.6654). For the Little finger (not shown on Figure 2), the results revealed a monotonic *decrease* in CSE from GoCue to Late RT. Namely, CSE decreased at Late RT (1.572 ± 0.253 mV) as compared to both Early RT (1.764 ± 0.278 mV; β = 0.1929 ± 0.0867; *p* = 0.0261) and GoCue (1.975 ± 0.330 mV; β = 0.4035 ± 0.1183; *p* = 0.0006). CSE also decreased from GoCue to Early RT (β = 0.2106 ± 0.0967; *p* = 0.0294). Overall, these results show that CSE increased in a digit-*specific* manner during movement preparation in the ICF experiment.

**Figure 2.**
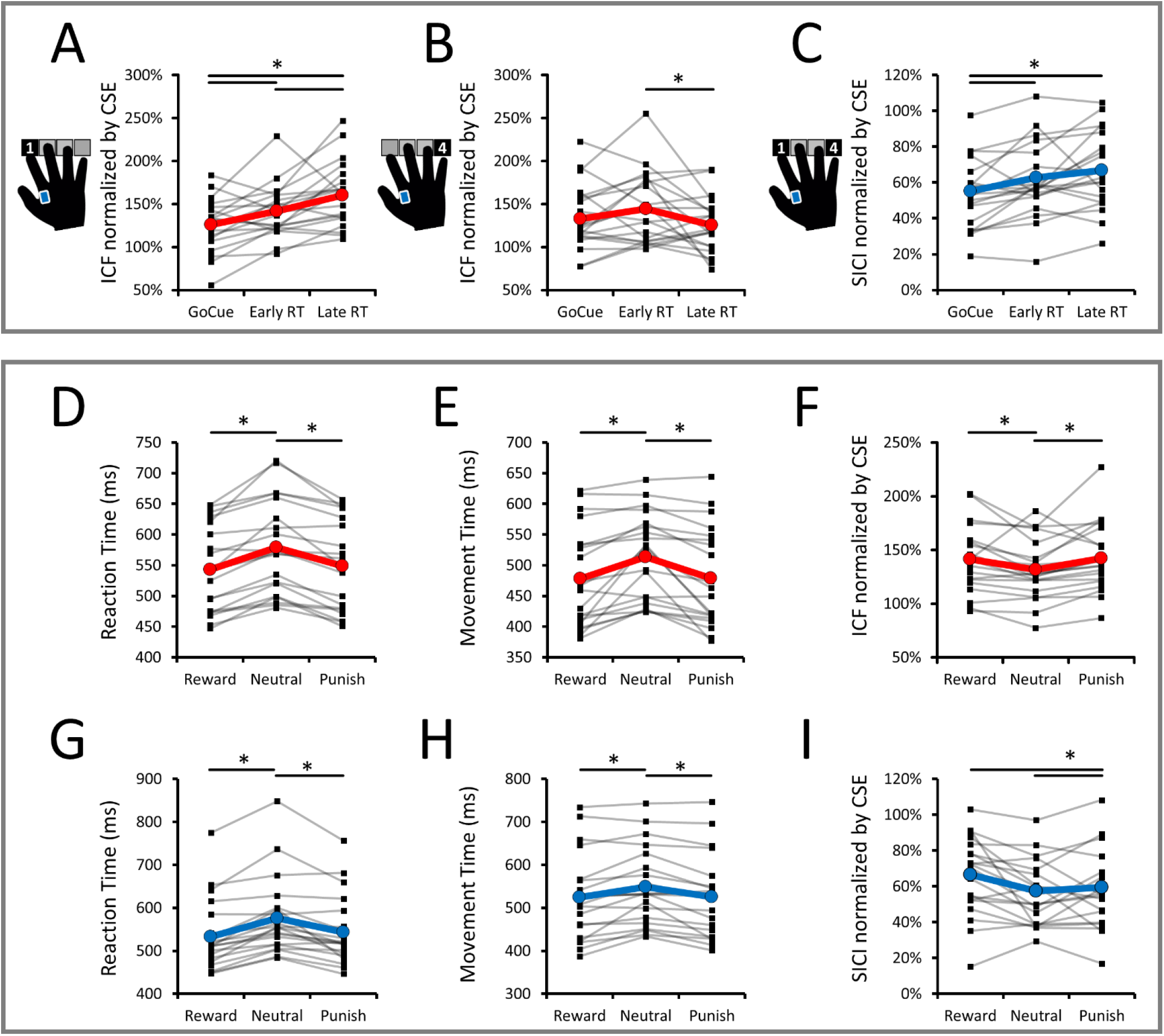
ICF and SICI Changes During Movement Preparation. **(A)** *ICF changes in the FDI muscle when the index finger initiated the sequence*. **(B)** *ICF changes when the little finger initiated the sequence*. Nearing movement onset increased ICF in the FDI muscle, but only when the index finger initiated the sequence. This suggests an effector-specific increase in M1’s GLUTergic facilitation during preparation. **(C)** *SICI changes in the FDI muscles when either the index or little fingers initiated the sequence*. SICI changes did not interact with the initiating finger, suggesting a broad effector-unspecific decrease in GABA_A_ inhibition during preparation. **(D)** *ICF Experiment – RT data per level of Incentives*. **(E)** *ICF Experiment – MT data per level of Incentives*. ***(F)*** *ICF Experiment – ICF data per level of Incentives*. Whilst panels (D) and (E) confirm that incentives enhanced performance, panel (F) shows that both rewards and punishments increased ICF as compared to the neutral condition. The increased ICF did not interact with the Time Points or Initiating Finger factors, suggesting that incentives enhance performance by increasing GLUTergic activity in a temporal- and effector-*unspecific* manner during movement preparation. **(G)** *SICI Experiment – RT data per level of Incentives*. **(H)** *SICI Experiment – MT data per level of Incentives*. **(I)** *SICI Experiment – SICI data per level of Incentives*. Whilst panels (G) and (H) confirm that incentives enhanced performance, panel (I) shows that rewards decreased SICI as compared to both neutral and punishments conditions. The decreased SICI also did not interact with the Time Points and Initiating Finger factors, suggesting that rewards further contributed to invigorating performance by decreasing GABA_A_ inhibition in a temporal- and effector-*unspecific* manner during preparation. For panels (D) to (I), “Punish” means “Punishment”. For all panels, individual data (n = 20) with their respective means are shown^38^.

For the SICI experiment, the results revealed an effect of Time Points (χ^2^ = 18.009; *p* = 0.0001), an effect of Initiating Finger (χ^2^ = 4.399; *p* = 0.0360), and a three-way Incentives * Time Points * Initiating Finger interaction (χ^2^ = 10.014; *p* = 0.0402). The effect of Time Points (not shown on Figure 2) revealed greater CSE at Late RT (1.966 ± 0.233 mV) as compared to both GoCue (1.616 ± 0.224 mV; β = 0.822 ± 0.058; *p* = 0.0083) and Early RT (1.510 ± 0.196 mV; β = 1.302 ± 0.081; *p* < 0.0001). CSE at Early RT and GoCue did not significantly differ (β = 1.071 ± 0.053; *p* = 0.1682). The effect of Initiating Finger (not shown on Figure 2) revealed greater CSE when the Index (1.787 ± 0.231 mV) initiated the sequences as compared to the Little finger (1.591 ± 0.199 mV; β = 1.123 ± 0.062; *p* = 0.0360). The three-way interaction was decomposed by conducting simple effects of Time Points separately for each level of Incentives and Initiating Finger. Overall, the results revealed main effects of Time Points selectively when the Index initiated the sequences, regardless of the level of Incentives (not shown on Figure 2). Specifically, when the Index initiated the sequences, there were simple effects of Time Points in the Reward (χ^2^ = 10.470; *p* = 0.0053), Neutral (χ^2^= 8.908; *p* = 0.0116) and Punishment conditions (χ^2^ = 12.580; *p* = 0.0019). Oppositely, when the Little finger initiated the sequences, there were no simple effects of Time Points in the Reward (χ^2^ = 3.368; *p* = 0.2784), Neutral (χ^2^ = 2.144; *p* = 0.3422), and Punishment conditions (χ^2^ = 4.512; *p* = 0.3144). Overall, these results show that CSE increased in a digit-*specific* manner during movement preparation in the SICI experiment.

### Digit-*specific* ICF increases and digit-*unspecific* SICI decreases during preparation

As above, generalised mixed linear models with the same fixed effects as above were conducted to investigate changes in normalised ICF and SICI data during movement preparation. Only the effects and interactions involving Time Points and Initiating Finger are reported in this section. For the ICF experiment (Figure 2A and 2B), the results revealed a Time Points * Initiating Finger interaction (χ^2^ = 42.174; *p* < 0.0001). A breakdown of this interaction revealed simple effects of Time Points when the Index (Figure 2A; χ^2^ = 13.104; *p* = 0.0028), but not when the Little finger (Figure 2B; χ^2^ = 4.786; *p* = 0.0914), initiated the sequence. For the Index finger (Figure 2A), ICF was greater at both Late RT (162 ± 8%; β = 0.3333 ± 0.0931; *p* = 0.0003) and Early RT (145 ± 9%; β = 0.1659 ± 0.0727; *p* = 0.0225) as compared to GoCue (128 ± 8%). ICF was also greater at Late RT than at Early RT (β = 0.1674 ± 0.0839; *p* = 0.0459). Overall, these results show that ICF monotonically increased during movement preparation in a digit-*specific* manner.

For the SICI experiment (Figure 2C), the results revealed an effect of Time Points (χ^2^ = 20.023; *p* < 0.0001), but no effect of Initiating Finger (χ^2^ = 1.168; *p* = 0.2798), no Time Points * Initiating Finger interaction (χ^2^ = 1.552; *p* = 0.4602), and no three-way Incentives * Time Points * Initiating Finger interaction (χ^2^ = 7.554; *p* = 0.1094). Breakdown of the effect of Time Points revealed reduced SICI (greater MEP amplitude) at both Late RT (59 ± 5%; β = 0.8424 ± 0.0372; *p* = 0.0001) and Early RT (57 ± 5%; β = 0.8727 ± 0.0437; *p* = 0.0098) as compared to GoCue (50 ± 4%; Figure 2C). SICI measured at Late and Early RT did not significantly differ (β = 1.0359 ± 0.0644; *p* = 0.5700). Overall, these results show that SICI decreases during movement preparation in a digit-*unspecific* manner.

### Incentives enhanced behavioural performance, irrespective of whether TMS pulses were delivered

To confirm that rewards and punishments enhanced performance as compared to the neutral condition, RT and MT data were compared across Incentives (Reward, Neutral, Punishment) and TMS Trial Types (ICF or SICI, CSE, No TMS). Overall, the results show that TMS pulses disrupted performance by slowing RTs and MTs and – more importantly – incentives robustly enhanced performance across all TMS Trial Types. In addition, the faster RTs and MTs within Reward and Punishment conditions were not accompanied by decreases in sequence accuracy (success rates).

For the ICF Experiment (Figure 2D and 2E), results from the RT data showed an effect of Incentives (χ^2^ = 213.402, *p* < 0.0001), an effect of TMS Trial Type (χ^2^ = 11.987, *p* = 0.0025), but no interaction between the two factors (χ^2^ = 1.455, *p* = 0.8345). For Incentives (Figure 2D), RTs were faster in both the Reward (552 ± 20ms; β = -0.0351 ± 0.0026; *p* < 0.0001) and Punishment conditions (556 ± 20ms; β = -0.0307 ± 0.0026; *p* < 0.0001) as compared to the Neutral one (587 ± 20ms). RTs did not significantly differ between the Reward and Punishment conditions (β = -0.0043 ± 0.0025; *p* = 0.0813). For TMS Trial Type (not shown on Figure 2), delivering ICF (585 ± 22ms) slowed RT as compared to both CSE (561 ± 20ms; β = 0.0243 ± 0.0078; *p* = 0.0027) and NoTMS trials (550 ± 19ms; β = 0.0354 ± 0.0105; *p* = 0.0012). Delivering CSE did not slow RT as compared to NoTMS trials (β = 0.0110 ± 0.0069; *p* = 0.0980). Overall, this shows that delivering ICF pulses during movement preparation slowed RT, but reward and punishment nonetheless successfully enhanced RT data across all TMS Trial Types. For MT data (Figure 2E), there was an effect of Incentives (χ^2^ = 10.630; *p* = 0.0049), TMS Trial Type (χ^2^ = 93.770; *p* < 0.0001), and an interaction between the two factors (χ^2^ = 11.830, *p* = 0.0187). For Incentives, MTs were faster in both Reward (486 ± 21ms; β = -0.0346 ± 0.0107; *p* = 0.0018) and Punishment conditions (486 ± 21ms; β = -0.0341 ± 0.0107; *p* = 0.0022) as compared to the Neutral one (520 ± 19ms). MTs on Reward and Punishment conditions did not significantly differ (β = 0.0006 ± 0.0033; *p* = 0.8651). The effect of TMS Trial Type (not shown on Figure 2) revealed that delivering ICF (511 ± 20ms) slowed MT as compared to both CSE (501 ± 20ms; β = 0.0101 ± 0.0029; *p* = 0.0004) and NoTMS trials (481 ± 20ms; β = 0.0300 ± 0.0031; *p* < 0.0001). Delivering CSE also slowed MT as compared to NoTMS trials (β = 0.0200 ± 0.0031; *p* < 0.0001). Breakdown of the Incentives * TMS Trial Type interaction revealed simple effects of Incentives for NoTMS (χ^2^ = 11.436; *p* = 0.0033), CSE (χ^2^ = 14.174; *p* = 0.0008), and ICF trials (χ^2^ = 6.142; *p* = 0.0464), confirming that rewards and punishments enhanced MT across all TMS trial types. Overall, this shows that TMS pulses delivered during preparation slowed MT, but also that incentives nonetheless successfully enhanced MT data across all TMS Trial Types.

For the SICI experiment (Figure 2G and 2H), the RT data showed an effect of Incentives (χ^2^ = 30.249, *p* < 0.0001), an effect of TMS Trial Type (χ^2^ = 12.627, *p* = 0.0018), but no interaction between the two factors (χ^2^ = 1.812, *p* = 0.7703). For Incentives (Figure 2G), RTs were faster in both the Reward (546 ± 24ms; β = -0.0425 ± 0.0084; *p* < 0.0001) and Punishment conditions (555 ± 23ms; β = -0.0336 ± 0.0105; *p* = 0.0019) as compared to the Neutral one (588 ± 26ms). RTs did not significantly differ between the Reward and Punishment conditions (β = -0.0089 ± 0.0052; *p* = 0.0903). For TMS Trial Type (not shown on Figure 2), CSE slowed RT (581 ± 27ms) as compared to SICI (559 ± 24ms; β = 0.0214 ± 0.0060; *p* = 0.0006) and NoTMS trials (550 ± 21ms; β = -0.0303 ± 0.0099; *p* = 0.0033). RTs on NoTMS and SICI trials did not significantly differ (β = -0.0089 ± 0.0061; *p* = 0.1461). Overall, this shows that delivering TMS pulses during preparation slowed RT, but incentives nonetheless successfully enhanced RT data across all TMS Trial Types. For MT data (Figure 2H), there was an effect of Incentives (χ^2^ = 9.752; *p* = 0.0076) and TMS Trial Type (χ^2^ = 114.934; *p* < 0.0001), but no interaction between the two factors (χ^2^ = 0.911; *p* = 0.9230). For Incentives (Figure 2H), MTs were faster in both Reward (535 ± 27ms; β = -0.0225 ± 0.0082; *p* = 0.0092) and Punishment conditions (535 ± 26ms; β = -0.0225 ± 0.0072; *p* = 0.0027) as compared to the Neutral one (558 ± 24ms). MTs on Reward and Punishment conditions did not significantly differ (β < -0.0001 ± 0.0035; *p* = 0.9927). The effect of TMS Trial Type (not shown on Figure 2) revealed slower MTs when CSE (559 ± 25ms; β = -0.0330 ± 0.0031; *p* < 0.0001) and SICI were delivered (542 ± 25ms; β = -0.0153 ± 0.0031; *p* < 0.0001) as compared to NoTMS trials (526 ± 25ms). Delivering CSE also slowed MT as compared to SICI trials (β = 0.0177 ± 0.0028; *p* < 0.0001). This shows TMS pulses delivered during preparation slowed MT, but that incentives nonetheless successfully enhanced MT data across all TMS Trial Types.

Further analyses were conducted to determine if the improvements in RTs and MTs were accompanied by changes in accuracy (success rates) at executing the sequences. Since this variable is binary (on a trial-per-trial basis), generalised mixed linear models with a binomial distribution were conducted. For the ICF Experiment, the results revealed an effect of Incentives (χ^2^ = 11.683, *p* = 0.0029), an effect of TMS Trial Type (χ^2^ = 54.787, *p* < 0.0001), but no interaction between the two factors (χ^2^ = 7.346, *p* = 0.1187). For Incentives (not shown on Figure 2), results revealed that accuracy was higher in the Reward condition (92.3 ± 1.2%) as compared to the Neutral one (89.7 ± 1.6; ^β = 0.7330 ± 0.0671; *p* = 0.0021). The Punishment condition (91.2 ± 1.4%) did not significantly differ from the Reward (^β = 0.8678 ± 0.0832; *p* = 0.1389) and Neutral conditions (^β = 1.1839 ± 0.1072; *p* = 0.0936). For TMS Trial Type (not shown on Figure 2), results revealed that accuracy was higher on both NoTMS (93.3 ± 1.1%; ^β = 1.9434 ± 01878; *p* < 0.0001) and CSE trials (91.5 ± 1.3%; ^β = 0.6710 ± 0.0533; *p* < 0.0011) as compared to ICF ones (87.8 ± 1.8%). Error rates on NoTMS trials were also lower than on CSE trials (^β = 1.3041 ± 0.1313; *p* = 0.0084). Overall, these results show that Incentives did not alter accuracy (success rates) and that it was the highest on NoTMS trials.

For the SICI Experiment, the results revealed an effect of TMS Trial Type (χ^2^ = 66.248, *p* < 0.0001), but no effect of Incentives (χ^2^ = 0.107, *p* = 0.9479), and no interaction between the two factors (χ^2^ = 7.210, *p* = 0.1252). For TMS Trial Type (not shown on Figure 2), results revealed that accuracy was greater on CSE trials (86.2 ± 2.2%) as compared to both NoTMS (92.3 ± 1.3%; ^β = 1.912 ± 0.1735; *p* < 0.0001) and SICI ones (90.9 ± 1.5%; ^β = 0.6233 ± 0.0476; *p* < 0.0001). Accuracy did not significantly differ between NoTMS and SICI trials (^β = 1.1919 ± 0.1138; *p* = 0.0658). Overall, these results show that Incentives did not alter accuracy (success rates) and that it was the highest on NoTMS trials.

### Incentives altered CSE in the SICI experiment, but not in the ICF one

These analyses assessed if Incentives altered CSE during preparation. For the ICF experiment (not shown on Figure 2), the results revealed no effect of Incentives (χ^2^ = 4.019; *p* = 0.1340), no Incentives * Time Points interaction (χ^2^ = 5.903; *p* = 0.2065), no Incentives * Initiating Finger (χ^2^ = 2.167; *p* = 0.3383), and no three-way interaction (χ^2^ = 3.931; *p* = 0.4154). For the SICI experiment, the results revealed an effect of Incentives (χ^2^ = 6.469; *p* = 0.0394), but no Time Points * Incentives interaction (χ^2^ = 5.831; *p* = 0.2122), and no Incentives * Initiating Finger interaction (χ^2^ = 0.621; *p* = 0.7329). The effect of Incentives (not shown on Figure 2) revealed that Punishment (1.764 ± 0.221 mV) increased CSE as compared to both the Reward (1.643 ± 0.206 mV; β = 1.0740 ± 0.0330; *p* = 0.0306) and Neutral conditions (1.655 ± 0.208 mV; β = 1.0660 ± 0.0328; *p* = 0.0576). The Reward and Neutral conditions did not significantly differ (β = 1.0008 ± 0.0309; *p* = 0.8055). Overall, this shows that rewards and punishments did not alter CSE in the ICF experiment, but that punishments increased CSE as compared to both reward and neutral conditions in the SICI experiment.

### Both rewards and punishments increased ICF, but only rewards decreased SICI

These analyses assessed if Incentives altered ICF and SICI during preparation. For the ICF experiment (Figure 2F), there was an effect of Incentives (χ^2^ = 9.407; *p* = 0.0091), but no Incentives * Time Points (χ^2^ = 2.741; *p* = 0.6020), no Incentives * Initiating Finger (χ^2^ = 1.321; *p* = 0.5166) and no three-way interaction (χ^2^ = 6.742; *p* = 0.1502). Breakdown of the effect of Incentives (Figure 2F) revealed greater ICF in both the Reward (146 ± 7%; β = 0.1251 ± 0.0446, *p* = 0.0075) and Punishment conditions (145 ± 7%; β = 0.1202 ± 0.0447, *p* = 0.0108) as compared to the Neutral one (133 ± 6%). ICF in Reward and Punishment conditions did not significantly differ (β = 0.0049 ± 0.0396, *p* = 0.9022). Overall, this shows that both incentives increased ICF as compared to the neutral condition. This also shows that the effects of incentives on ICF did not statistically interact with Time Points or the Initiating Finger, suggesting that incentives enhance performance by increasing GLUTergic activity in a temporal- and effector-*unspecific* manner during movement preparation.

For the SICI experiment (Figure 2I), there was an effect of Incentives (χ^2^ = 15.735; *p* = 0.0091), but no Incentives * Time Points (χ^2^ = 6.006; *p* = 0.1987), no Incentives * Initiating Finger (χ^2^ = 2.050; *p* = 0.3589), and no three-way interaction (χ^2^ = 7.554; *p* = 0.1094). Breakdown of the effect of Incentives (Figure 2I) revealed decreased SICI in the Reward condition (60 ± 5%) as compared to both the Punishment (53 ± 4%; β = 0.8794 ± 0.0343; *p* = 0.0015) and Neutral ones (52 ± 4%; β = 0.0871 ± 0.0340; *p* = 0.0006). SICI on Neutral and Punishment conditions did not significantly differ (β = 1.0100 ± 0.0395; *p* = 0.7990). Overall, this shows that rewards decreased SICI as compared to both punishment and neutral conditions. This also shows that the effect of reward on SICI did not statistically interact with Time Points or the Initiating Finger, suggesting that rewards enhance performance by decreasing GABA_A_-mediated inhibition in a temporal- and effector-*unspecific* manner during movement preparation.

### RATIONAL OF THE FEEDBACK PROCESSING SESSION

The above results suggest that, as compared to the neutral condition, both rewards and punishments increased GLUTergic facilitation, whereas only rewards decreased GABA_A_ inhibition during preparation. Interestingly, the results also revealed that these effects of Incentives on ICF and SICI were temporally- and effector-*unspecific*, as they did not interact with the digit-*specific* ICF increases and digit-*unspecific* SICI decreases observed during preparation as movement onset approached. This suggests that incentive-induced ICF and SICI changes reflect temporally unspecific mechanisms by which rewards and punishments alter M1’s intracortical excitability to enhance performance, as previous TMS work would suggest^40–42^. If valid, this suggests that incentives enhance performance not by solely altering movement preparation, but by inducing a tonic brain state that permeates beyond movement preparation. The objective of this additional session was to evaluate this possibility.

Here, the effect of reward and punishment feedback delivery (> 1,000ms after the execution of the sequences) on ICF and SICI was evaluated. The same experimental procedures as in Figure 1A and 1B were used except for the time points at which TMS pulses were delivered (Feedback [FB] Onset, 500ms, and 1,000ms after; Figure 3A). Hereafter, this is referred to as the Feedback Processing session. If ICF and SICI during feedback processing are modulated similarly as in movement preparation, this would suggest that increased GLUTergic and decreased GABA_A_ inhibition are (tonic) temporally-*unspecific* mechanisms by which incentives enhance motor performance. Conversely, if ICF and SICI are not altered during feedback processing, this would suggest that rewards and punishments enhance motor performance by acutely altering ICF and SICI during movement preparation only. Importantly, to strengthen the possibility that incentives could selectively alter movement preparation but not feedback processing, the same groups of 20 participants that showed incentive-*specific* ICF and SICI changes during movement preparation were recruited for the Feedback Processing session.

**Figure 3.**
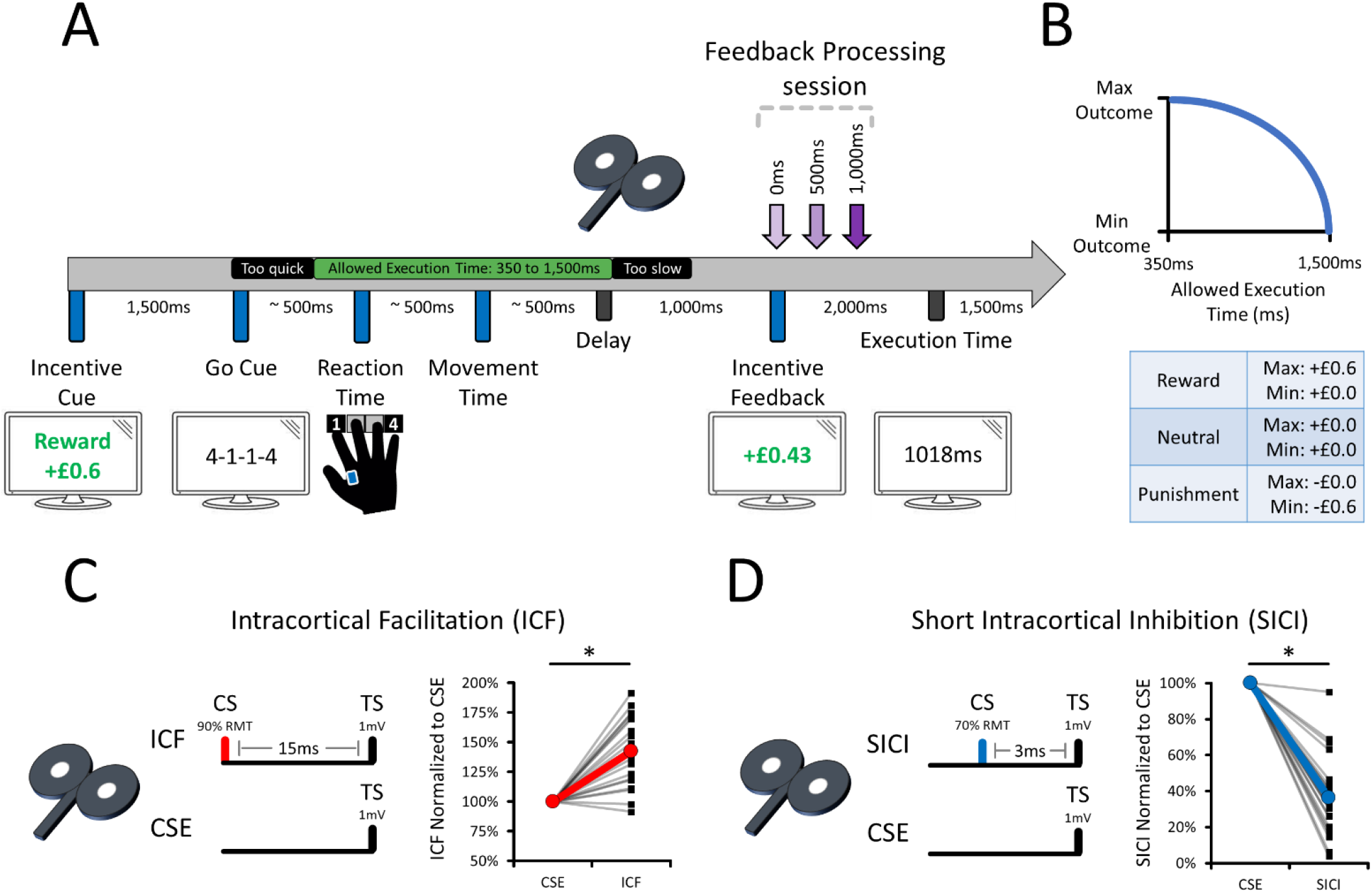
Feedback Processing session for both the ICF and SICI Experiments. **(A)** *Chronology of a typical trial*. For both Experiments, the same procedures as in Figure 1 were used, except for the time points at which TMS pulses were delivered; TMS pulses were delivered either at Incentive Feedback onset (FB Onset, or 0ms), 500ms or 1,000ms later to investigate if incentives alter ICF and SICI during feedback processing. EMG data were also recorded from the FDI muscle. **(B)** *The decay function used to adjust the incentives based on performance*. “Max” and “Min” refer to the maximal and minimal possible outcome on a given trial. Allowed execution time is defined as the sum of reaction time and movement time (as depicted in panel A). **(C)** *TMS parameters for the ICF Experiment (n = 20)* successfully induced ICF at rest, as assessed by delivering 30 CSE and 30 ICF pulses. **(D)** *TMS parameters for the SICI Experiment (n = 20)* successfully induced SICI at rest, as assessed by delivering 30 CSE and 30 SICI pulses. For panels (C) and (D), individual data with their respective means are shown^38^. RMT means resting motor threshold.

## RESULTS – FEEDBACK PROCESSING SESSION

### ICF and SICI were reliably observed at rest

The Feedback Processing session was conducted using the same TMS parameters as in the Movement Preparation session (Figure 3C and 3D). The results revealed significant M1 facilitation with ICF (Figure 3C; 1.389 ± 0.167 mV) as compared to CSE (0.988 ± 0.123 mV; β = 0.4006 ± 0.0740; *p* < 0.0001). In addition, there was significant M1 inhibition with SICI (Figure 3D; 0.379 ± 0.067 mV) as compared to CSE (1.089 ± 0.143 mV; β = 0.7095 ± 0.1015; *p* < 0.0001). Note that Figure 3C and 3D reports ICF and SICI values normalised (%) to the average of all CSE trials within each participant. Overall, this confirms that reliable ICF and SICI was observed at rest across participants, replicating results from the Movement Preparation session.

### Incentives enhanced motor performance

The next set of analyses sought to confirm that rewards and punishments enhanced motor performance as compared to the neutral condition. Here, since TMS pulses were delivered >1,000ms after movement completion^41,42^, the effects of TMS Trial Type and Initiating Finger were not included in the analyses. Overall, the results show that incentives successfully enhanced performance without decreasing accuracy (success rates), replicating the behavioural results from the Movement Preparation session.

For the ICF experiment (Figure 4A and 4B), the RT data (Figure 4A) showed an effect of Incentives (χ^2^ = 13.740; *p* = 0.0010), which was driven by faster RTs on both the Reward (536 ± 15ms; β = -0.0392 ± 0.0108; *p* = 0.0004) and Punishment conditions (540 ± 15ms; β = -0.0347 ± 0.0094; *p* = 0.0003) as compared to the Neutral one (575 ± 18ms). RTs on Reward and Punishment conditions did not significantly differ (β = -0.0045 ± 0.0034; *p* = 0.1848). For MT data (Figure 4B), there was an effect of Incentives (χ^2^ = 7.837; *p* = 0.0199). MTs were faster for both the Reward (446 ± 19ms; β = -0.0350 ± 0.0128; *p* = 0.0095) and Punishment conditions (449 ± 19ms; β = -0.0320 ± 0.0127; *p* = 0.0176) as compared to the Neutral one (481 ± 18ms). MTs on Reward and Punishment conditions did not significantly differ (β = 0.0030 ± 0.0029; *p* = 0.3038). Overall, this confirms that incentives enhanced motor performance in the ICF experiment.

**Figure 4.**
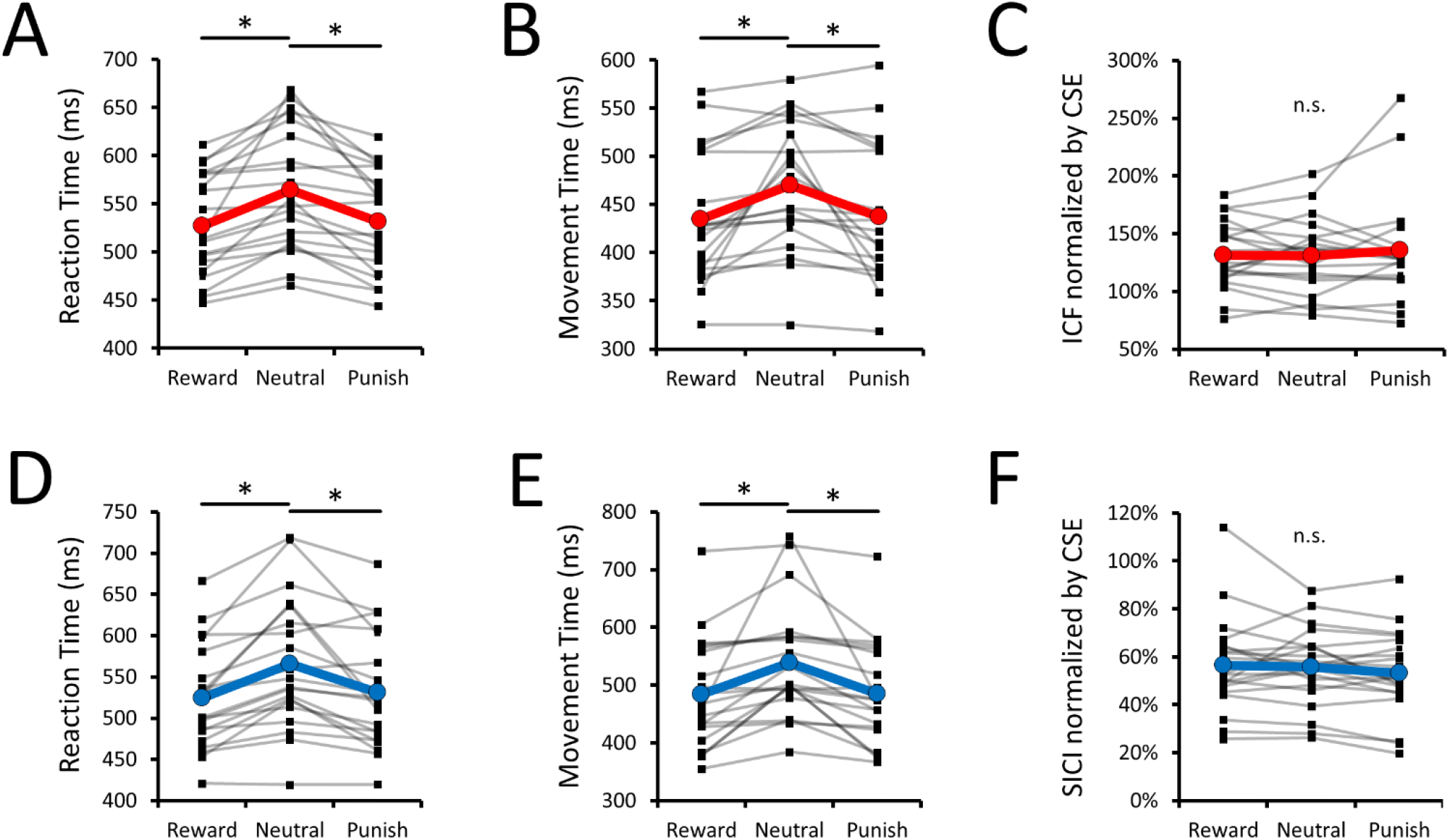
ICF and SICI Changes During Feedback Processing. **(A)** *ICF Experiment – RT data per level of Incentives*. **(B)** *ICF Experiment – MT data per level of Incentives*. ***(C)*** *ICF Experiment – ICF data per level of Incentives*. Whilst panels (A) and (B) confirm that incentives enhanced performance, panel (C) shows that both rewards and punishments failed to alter ICF as compared to the neutral condition. **(D)** *SICI Experiment RT data per level of Incentives*. **(E)** *SICI Experiment – MT data per level of Incentives*. **(F)** *SICI Experiment SICI data per level of Incentives*. Whilst panels (D) and (E) confirm that incentives enhanced performance, panel (F) shows that both rewards and punishments failed to alter SICI as compared to the neutral condition. Overall, this shows that incentives enhanced performance, but not by altering ICF or SICI upon incentive feedback delivery. As such, these results suggest that incentives selectively alter the processes of movement preparation to enhance motor performance. For all panels, “Punish” means “Punishment”. For all panels, individual data (n = 20) with their respective means are shown^38^.

For the SICI experiment (Figure 4D and 4E), the RT data (Figure 4D) showed an effect of Incentives (χ^2^ = 24.930; *p* < 0.0001). RTs were faster on both the Reward (534 ± 18ms; β = -0.0443 ± 0.0107; *p* < 0.0001) and Punishment conditions (542 ± 20ms; β = -0.0368 ± 0.0128; *p* = 0.0061) as compared to the Neutral one (579 ± 23ms). RTs on Reward and Punishment conditions did not significantly differ (β = 0.0076 ± 0.0048; *p* = 0.1141). For MT data (Figure 4E), there was an effect of Incentives (χ^2^ = 7.840; *p* = 0.0198). MTs were faster on both the Reward (498 ± 26ms; β = -0.0570 ± 0.0212; *p* = 0.0107) and Punishment conditions (500 ± 25ms; β = -0.0559 ± 0.0200; *p* = 0.0080) as compared to the Neutral one (555 ± 28ms). MTs on Reward and Punishment conditions did not significantly differ (β = 0.0011 ± 0.0041; *p* = 0.7768). Overall, this confirms that incentives enhanced motor performance in the SICI experiment.

Further analyses were conducted to determine if the improvements in RTs and MTs were accompanied by decreases in accuracy (success rates). Since this variable is binary (on a trial-per-trial basis), generalized mixed linear models with a binomial distribution were also conducted. For the ICF Experiment (not shown on Figure 4), the results revealed no effect of Incentives (χ^2^ = 1.697; *p* = 0.4281), showing that accuracy did not significantly differ between the Reward (93.8 ± 1.1%), Neutral (93.7 ± 1.1%) and Punishment conditions (94.4 ± 1.0%). For the SICI Experiment (not shown on Figure 4), the results revealed no effect of Incentives (χ^2^ = 4.046; *p* = 0.1323), showing that accuracy did not significantly differ between the Reward (95.1 ± 0.7%), Neutral (92.8 ± 1.4%) and Punishment conditions (94.2 ± 0.8%). Overall, this shows that accuracy (success rates) did not significantly differ amongst incentive conditions in both the ICF and SICI experiments.

### Incentives did not alter CSE in the ICF experiment, but punishments increased CSE in the SICI one

Generalised mixed linear models were conducted to evaluate CSE changes during feedback processing. The fixed effects included in the analyses were Incentives (Reward, Neutral, Punishment) and Time Points (FB onset, 500ms, 1,000ms). For the ICF experiment (not shown on Figure 4), the results revealed no effect of Incentives (χ^2^ = 3.490; *p* = 0.1747), no effect of Time Points (χ^2^ = 3.845; *p* = 0.1462), and no Incentives * Time Points interaction (χ^2^ = 1.714; *p* = 0.7882). This shows that CSE did not significantly differ between the Reward (1.944 ± 0.290 mV), Punishment (1.902 ± 0.304 mV), and Neutral (2.016 ± 0.304 mV) conditions.

For the SICI experiment (not shown on Figure 4), the results revealed an Incentives * Time Points interaction (χ^2^ = 22.973; *p* = 0.0001) and an effect of Time Points (χ^2^ = 17.670; *p* = 0.0001), but no effect of Incentives (χ^2^ = 3.225; *p* = 0.1994). The Incentives * Time Points interaction was decomposed by conducting simple effects of Incentives at each level of Time Points (not shown on Figure 4). At FB Onset, despite results revealing a simple effect of Incentives (χ^2^ = 5.762; *p* = 0.0561), CSE did not significantly differ in the Reward (1.963 ± 0.104 mV; β = 0.0044 ± 0.0546; *p* = 0.9361) and Punishment conditions (1.869 ± 0.105 mV; β = -0.0894 ± 0.0423; *p* = 0.1032) as compared to the Neutral one (1.959 ± 0.106 mV). CSE on Reward and Punishment conditions also did not significantly differ (β = -0.0938 ± 0.0522; *p* = 0.1086). At 500ms, the simple effect of Incentives (χ^2^ = 8.206; *p* = 0.0165) revealed greater CSE in Punishment (1.923 ± 0.105 mV) as compared to the Reward condition (1.782 ± 0.102 mV; β = 0.1407 ± 0.0491; *p* = 0.0126). Both the Reward (β = 0.0926 ± 0.0484; *p* = 0.0837) and Punishments conditions (β = 0.0482 ± 0.0407; *p* = 0.2365) did not significantly differ from the Neutral one (1.875 ± 0.105 mV). At 1000ms, the simple effect of Incentives (χ^2^ = 10.796; *p* = 0.0045) revealed greater CSE in Punishment (1.910 ± 0.105 mV) as compared to both the Reward (1.758 ± 0.102 mV; β = 0.1527 ± 0.0484; *p* = 0.0024) and Neutral conditions (1.824 ± 0.104 mV; β = 0.0861 ± 0.0384; *p* = 0.0378). The Reward and Neutral conditions did not significantly differ (β = 0.0666 ± 0.0464; *p* = 0.1507). Overall, this shows that CSE did not significantly differ amongst incentives at FB onset, that CSE increased in Punishment as compared to the Reward condition at 500ms, and that CSE remained increased in Punishment as compared to both Reward and Neutral conditions at 1000ms.

### Incentives did not alter ICF or SICI during feedback processing

The same analyses as above were conducted on ICF and SICI data. For the ICF experiment (Figure 4C), the results revealed no effect of Incentives (χ^2^ = 0.859; *p* = 0.6508), no effect of Time Points (χ^2^ = 1.511; *p* = 0.4699), and no Incentives * Time Points interaction (χ^2^ = 5.508; *p* = 0.2391). Overall, this shows that ICF did not significantly differ amongst the Reward (135 ± 7%), Neutral (135 ± 8%) and Punishment conditions (142 ± 11%). For the SICI experiment (Figure 4F), the results revealed no effect of Incentives (χ^2^ = 3.342; *p* = 0.1881), no effect of Time Points (χ^2^ = 0.898; *p* = 0.6382), and no Incentives * Time Points interaction (χ^2^ = 3.077; *p* = 0.5451). Overall, this shows that SICI did not significantly differ amongst the Reward (57 ± 4%), Neutral (56 ± 4%) and Punishment conditions (53 ± 4%).

## DISCUSSION

This work investigated if rewards and punishments enhance motor performance through circuit-specific changes in GLUTergic facilitation and GABA_A_-mediated inhibition in M1 during movement preparation, as assessed with ppTMS. First, the results revealed digit-*specific* CSE and ICF increases and digit-*unspecific* SICI decreases as movement onset approached during preparation. These findings suggest that M1 increases CSE and GLUTergic activity in an effector-*specific* manner, whilst broadly decreasing local GABA_A_ inhibition in an effector-*unspecific* manner during movement preparation. Second, behaviourally, rewards and punishments successfully quickened RTs and MTS as compared to the neutral conditions in both sessions and experiments, suggesting that incentives systematically enhanced motor performance. Importantly, both rewards and punishments increased ICF but only rewards decreased SICI during movement preparation, suggesting that incentives enhance performance through common (GLUTergic) and distinct (GABA_A_) circuit-specific changes in M1. Furthermore, this effect of incentives did not interact with the digit-*specific* ICF increases and digit-*unspecific* SICI decreases observed as movement onset approached, suggesting that rewards and punishments non-specifically enhance M1’s intracortical excitability during movement preparation. Finally, a control session investigating circuit-specific changes during Feedback Processing showed that ICF and SICI were not modulated post-movement upon (and following) the delivery of reward and punishment feedback. These differences suggest that incentives enhance performance by non-specifically altering ICF and SICI during movement preparation only. Globally, these results map the neurochemical mechanisms by which rewards and punishments enhance motor performance.

### Digit-*specific* CSE and ICF increases, but digit-*unspecific* SICI decreases during preparation

An important result of this study is that CSE increased during preparation in the digit that would initiate the cued finger-press sequence, as CSE increases were only observed in the EMG data of the FDI muscle when the index finger initiated the 4-element finger-press sequences. These digit-*specific* CSE increases were observed in both the ICF and SICI experiments and replicate previous work (for a review, see ref ^43^). Here, two novel results are that M1 increased ICF as movement onset approached and that these ICF increases were also digit-*specific*, constituting the first human evidence indicating that M1 increases its intracortical GLUTergic activity in an effector-*specific* manner during preparation. Such ICF increases also suggest that changes in GLUTergic activity in M1 represent the effector-*specific* preparation and delivery of motor commands. This possibility is directly supported by anatomical work showing that GLUTergic pyramidal neurons located in M1’s intralaminar layer 5 send excitatory projections that activate spinal motor centers^44^, which animal work has shown to be responsible for the preparation and execution of motor commands^24,25^. Overall, this evidence suggests that the CSE increases observed during movement preparation are mediated by increased GLUTergic activity in M1, likely reflecting upcoming motor commands.

Another key result is that SICI decreased in M1 as movement onset approached, which is largely consistent with previous TMS work^21–23^ and suggests that M1 decreases its local GABA_A_-mediated inhibition during preparation. Importantly, the present results extend existing TMS work^21–23^ by showing for the first time that these SICI decreases were digit-*unspecific*; that is, SICI decreased in the EMG data of the FDI muscle regardless of whether the index or little finger initiated the 4-element finger-press sequence. Interestingly, CSE increased in a digit-*specific* manner during the SICI experiment, contrasting with the digit-*unspecific* SICI decreases. One possibility is that increases in CSE (and ICF) reflect upcoming (effector-*specific*) motor commands, whilst SICI decreases reflect the local state transition from preparation to execution in M1^45^, presumably explaining the lack of effector-specificity. This contention is compatible with the status quo hypothesis^46–48^, which posits that decreases in M1’s beta-band power (13-30 Hz) – reflecting decreases in M1’s GABA_A_-mediated inhibition^49–51^ – allow the current sensorimotor state to transition from preparation to execution. Anatomical evidence also supports that SICI changes should not be expected to effector-*specific*, as GABA_A_ interneurons have few^52^, if any^53^, corticospinal projections. Indeed, GABA_A_ interneurons are known to locally regulate network dynamics of projecting GLUTergic pyramidal neurons^52,54^, making them prime circuits to shape motor commands in M1’s GLUTergic neurons^25,55^ and/or to withhold premature motor responses^43,56^. Therefore, this evidence suggests that the present digit-*unspecific* SICI decreases during movement preparation do not reflect upcoming motor commands *per se*, but rather the locally regulated state transition from preparation to execution in M1.

### Rewards and punishments both increased ICF, but only rewards decreased SICI during preparation

Both rewards and punishments quickened RTs and MTs without decreases in accuracy (success rates) across both the ICF and SICI experiments. These motor performance enhancements were robust to the disruptive effects of TMS pulse delivery, confirming that incentives successfully enhanced motor performance.

Regarding the TMS data, rewards and punishments did not alter CSE as compared to the neutral condition during preparation in the ICF experiment. However, punishments were found to non-specifically increase CSE as compared to both rewards and neutral conditions in the SICI experiment. Whereas the former result contrasts with TMS work showing that rewarding stimuli increase CSE during preparation as compared to neutral ones^6,7,10,57^, the latter result aligns with other evidence showing that punishing stimuli increases CSE during preparation^8^. Although the exact reasons remain unclear, one possibility is that these previously reported CSE changes were driven by circuit-specific changes in M1, which the present study specifically investigated. Namely, here, a key novel result is that both rewards and punishments increased ICF during movement preparation as compared to the neutral condition, suggesting that incentives upregulate M1’s GLUTergic activity to enhance performance. Importantly, these ICF increases did not interact with the digit-*specific* ICF increases observed as movement onset approached, suggesting that rewards and punishments increase GLUTergic activity in a non-specific manner during movement preparation. These results are largely consistent with animal work that shows that the VTA quickly regulates M1’s excitability^26^ by increasing GLUTergic activity throughout the entire M1^27^, suggesting that the present non-specific ICF increases in M1 have a subcortical origin. Furthermore, both incentives increased ICF, suggesting that stimuli with positive (reward) and negative (punishment) motivational value recruit a common neurochemical pathway in M1 to guide behaviours (see refs ^4,58^ for further support). Therefore, this evidence indicates that both rewards and punishments non-specifically increase M1’s intracortical GLUTergic activity during movement preparation to enhance motor performance.

In contrast to ICF, only rewards decreased SICI as compared to both punishment and neutral conditions during preparation, suggesting that rewards induce additional circuit-specific changes in M1 as compared to punishments. In a manner similar to ICF changes, the reward-specific SICI decreases did not interact with the digit-*unspecific* SICI decreases observed as movement onset approached, also suggesting non-specific effects of rewards on M1’s GABA_A_-mediated inhibition during preparation. Assuming that reward-induced motor performance enhancements are driven by enhanced dopaminergic activity (although see ref ^59^), one possibility is that the present SICI decreases were driven by dopaminergic pathways. In support, VTA dopaminergic projections have been shown to enhance M1’s excitability and crucially contribute to motor performance and learning (see ref ^28^ for a review), which could account for the present reward-specific SICI decreases. However, other evidence indicates that enhancing dopaminergic activity should rather increase SICI. Namely, activating M1’s D2-like receptors using quinpirole has been shown to increase the excitability of M1’s putative GABA_A_ interneurons^30^ and human TMS studies have shown that administration of dopamine agonists increases SICI (see ref ^14^ for a review). This evidence shows that increasing dopaminergic activity in M1 increases GABA_A_-mediated inhibition and SICI, which is incompatible with the possibility that rewards decrease SICI by increasing dopaminergic activity in M1. An alternative possibility is that increased noradrenergic activity accounts for the present SICI result. Namely, M1 receives considerable noradrenergic innervation^60^, and increases in noradrenergic activity are known to enhance performance of goal-directed behaviours^61,62^ as well as decrease SICI^14^, suggesting that increased noradrenergic activity accounts for the reward-specific SICI decreases. As a result, a possible relationship between SICI and dopaminergic or noradrenergic remains unclear. Nonetheless, one viable interpretation is that rewards enhance performance by decreasing SICI during preparation to facilitate the local transition from preparation to execution in M1 (as per the status quo hypothesis^46–48^). As GABA_A_-mediated inhibition is increasingly regarded as a key contributor to learning and memory consolidation^52,63^, the selective effects of rewards on SICI may also explain the added value of rewards on motor learning and memory consolidation as compared to punishments^64–69^.

It should be noted that delivering TMS pulses over the left M1 disrupted motor performance in the right hand by slowing RTs and MTs, obscuring the possibility to demonstrate an unbiased relationship between TMS-measured ICF, SICI, or CSE and changes in motor performance in the present work. However, as the present results suggest that rewards and punishments altered ICF and SICI in a manner that was effector-unspecific, future work could relate MEPs from the (disrupted) non-responding effector (e.g., left hand) with motor performance enhancements measured from the (undisrupted) responding effector (e.g., right hand). This would offer an unbiased approach to associate TMS-measured variables to changes in motor performance.

### Incentives did not alter ICF or SICI during feedback processing

The results from the feedback processing session revealed that rewards and punishments enhanced motor performance in both the ICF and SICI experiments. The results also revealed that CSE was not altered in the ICF experiment, but that punishments non-specifically enhanced CSE as compared to rewards in the SICI experiment. Although it remains unclear why punishments would only increase CSE in the SICI but not the ICF experiments, such an increase in CSE as compared to a reward (but not neutral) condition is consistent with previous TMS work^42^. An important result from the Feedback Processing session is that rewards and punishments did not alter ICF or SICI when measured post-movement during incentive feedback processing. This suggests that rewards and punishments enhance performance by selectively altering M1’s GLUTergic and GABA_A_ intracortical circuits during movement preparation. This result is consistent with the shift of activity from unexpected reward outcomes to predictive reward cues observed in VTA neurons^70^, but contrasts with previous TMS studies showing that rewards increase SICI upon reward delivery^40,41^. However, in these studies, rewards were delivered randomly, suggesting that increases in SICI represented a prediction error (difference between expected and received outcome^71^) rather than the isolated effects of rewards on M1’s GABA_A_-mediated inhibition. Here, participants were provided incentive cues at the start of each trial and were *a priori* informed that incentives would be performance contingent. Thus, participants likely combined knowledge of the incentive cues with their performance self-evaluation to anticipate the upcoming incentive feedback, resulting in little or no prediction errors. Such an absence of prediction errors could explain the present absence of ICF and SICI changes upon incentive feedback. Overall, these results show that rewards and punishments enhance performance by increasing ICF and decreasing SICI during movement preparation only. One implication is that targeting M1’s GLUTergic and GABA_A_ signalling movement preparation – but not feedback processing – processes using pharmacological and/or non-invasive brain stimulation interventions can further enhance the neurorehabilitative potential of rewards in clinical settings^15–18^.

### Conclusion

The present results suggest that M1 increases its GLUTergic activity to reflect the preparation and delivery of effector-*specific* motor commands and that M1 also locally decreases its GABA_A_ inhibition in an effector-*unspecific* manner to transition from movement preparation to execution. The results also suggests that, to enhance motor performance, rewards and punishments both increase M1’s GLUTergic activity and that only rewards decrease GABA_A_ inhibition in a temporal- and effector-*unspecific* manner during movement preparation, but not during post-movement feedback processing. Collectively, these results map the neurochemical changes in M1 by which incentives enhance motor performance.,

## METHODS

### Participants

Two groups of 20 medication-free and self-reported neurologically healthy participants took part in the ICF (23.9 ± 0.7 years old; 9 females) and SICI experiments (24.5 ± 1.3 years old; 15 females). Participants were screened for TMS contraindications^72^ and provided their informed written consent (approved by the local institutional board; project # ERN_17-1541AP6). Participants were paid a minimum of £20, which was topped up with their performance-based earnings. Overall, participants earned an average (± SEM) of £35.03 ± 0.83. For each experiment, the same participants completed both the Movement Preparation and Feedback Processing sessions. The sessions were counterbalanced across participants to mitigate carry-over effects^73^ and, for a given participant, took place at the same time of day to control for circadian influences of excitability^74,75^. On average, 5.1 ± 1.8 days and 2.9 ± 0.8 days separated the Movement Preparation and Feedback Processing sessions of the ICF and SICI experiments, respectively. This resulted in a total of 80 sessions, each lasting approximately 2h30.

The neurophysiological processing of rewards and punishments can be influenced by participant’s individual sensitivity to incentives^76,77^. Here, for each participant, individual sensitivity to rewards and punishments was measured using the 20-item revised and clarified Sensitivity to Punishment and Sensitivity to Reward Questionnaire (SPSRQ-RC)^78^. Separate independent t-tests revealed no difference for Sensitivity to Punishment (t_(38)_ = 0.500; *p* = 0.6201; Cohen’s d = 0.158) and Sensitivity to Reward (t_(38)_ = 0.122; *p* = 0.9040; Cohen’s d = 0.038) between the two groups of 20 participants that took part in the ICF and SICI experiments. This suggests that participants showed homogenous individual sensitivity to incentives across the two experiments.

### Task, apparatus and timing

The task consisted of executing 4-element finger-press sequences as fast and accurately as possible following the onset of a visual GoCue (Figures 1A and 3A). To investigate motor performance enhancements, participants executed the 4-element finger-press sequences under Reward (max: +£0.6; min: +£0.0), Neutral (max: +£0.0; min: +£0.0), and Punishment conditions (max: +£0.0; min: -£0.6). The magnitude of rewards or punishments obtained were a decay function (Figure 1B and 3B) of participants’ total execution time (defined as the sum of RT and MT) on a given trial. The 4-element finger-press sequences were executed on a USB-wired keyboard (600 Microsoft ®). All visual stimuli were displayed on a 24-inch iiyama Prolite (B2409HDS) computer monitor (1920 × 1080 pixels; vertical refresh rate [55 -75Hz]). The behavioural experiments were run through custom-built scripts using MatLab (R2021b; MathWorks ®) and the PsychToolBox-3 interface. To calculate finger press timing (required to assess individual RT and MT), the PsychToolBox-3 function “KbQueueCheck” was used, resulting in submillisecond (> 1,000Hz) sampling of RTs and MTs. The 4-element finger-press sequences selectively comprised index and little finger presses, which were set as the “D” and “J” keyboard keys and labelled as digits “1” and “4”, respectively. Doing so resulted in 6 possible 4-element finger-press sequences: “1-1-4-4”, “1-4-1-4”, “1-4-4-1”, “4-4-1-1”, “4-1-4-1”, and “4-1-1-4”. The sequences were pseudorandomised into 24-trial cycles, each trial containing a single 4-element finger-press sequence. The trials (sequences) were equiprobable amongst Reward, Neutral, and Punishment conditions (see below). Trials were pseudorandomised such that the same 4-element finger-press sequence or Incentive condition (Reward, Neutral, Punishment) never repeated on adjacent trials.

Within a 24-trial cycle, the Reward (8 trials), Neutral (8 trials), and Punishment conditions (8 trials) each comprised 3 single pulses to evaluate CSE, 3 paired-pulse trials (either ICF or SICI), and 2 NoTMS trials. Each of the 3 single pulse and paired-pulse trials was delivered at different time points at or following GoCue onset (0ms, 200ms, and 400ms; Figure 1) or incentive feedback onset (0ms, 500ms, and 1,000ms; Figure 3; see below for a justification of the chosen time points). Each session consisted of a total of 432 trials (324 TMS and 108 NoTMS trials), which were separated into three Blocks of 144 trials (108 TMS and 36 NoTMS trials) each lasting ∼25min. Short breaks were provided to participants every 48 trials, to prevent the accumulation of fatigue.

A USB-wired Arduino Nano board with a Deek Robot Terminal Adapter was controlled through MatLab to externally trigger the TMS stimulators by sending rising edge 5V triggers. Based on 800 behavioural trials and bootstrapped estimations (100,000 samples), the Arduino hardware had a latency of ∼20 ± 3ms (± SEM) between issuing the MatLab command the actual delivery of the 5V Trigger. This was offset by sending the 5V triggers 20ms earlier than the predefined time points in the Movement Preparation (0ms, 200ms and 400ms; Figure 1A) and Feedback Processing sessions (0ms, 500ms and 1,000ms; Figure 3A). Moreover, paired-pulse trials during the ICF and SICI experiments were triggered 15ms and 3ms earlier than the single pulse (CSE) trials, respectively. This was to account for the inter-pulse intervals on paired-pulse TMS trials (see below), allowing test stimuli to be delivered at a similar latency between single and paired-pulse trials.

### Typical trial chronology

An overview of the methods is provided in Figures 1 and 3 for the Movement Preparation and Feedback Processing sessions, respectively. A trial was initiated by displaying the Incentive Cue (“Reward+£0.6”, “Neutral +£0.0”, or “Punishment -£0.6”) for 1,500ms. To make the cues more salient, green-, grey-, and red-coloured fonts were used for the Reward, Neutral, and Punishment Incentive Cues, respectively. The GoCue was then displayed and consisted of the 4-element finger-press sequence to be executed. Participants were then allowed a total of 1,750ms to execute the sequence (“Allowed Execution Time” in Figures 1A and 3A, defined as the sum of RT and MT for the ongoing trial). Once the 4 keys were pressed or the 1,750ms limit was reached, whichever came first, the screen went black for a delay of 1,000ms. The Incentive Feedback was then displayed for 2,000ms, followed by the Execution Time of the ongoing trial for 1,500ms, after which the trial ended with a black screen. A fixed inter-trial interval of 1,500ms separated each trial. Each trial lasted ∼8,500ms, allowing sufficient time for the TMS stimulators to recharge between each trial.

### TMS time point delivery

To investigate intracortical excitability changes during movement preparation, TMS pulses were delivered at GoCue onset (0ms) as well as 200ms and 400ms following cue onset. These time points were chosen based on previous TMS work using similar timings to show reward-induced excitability changes during movement preparation^6,8,79^. They were also chosen to provide a measure excitability during the first (< 50%) and second halves (> 50%) of the RT period (similar to refs ^7,21,80^) whilst not exceeding the RT (> 100%), which was deemed critical to investigate the evolution of excitability during movement preparation. To investigate excitability changes during feedback processing, TMS pulses were delivered at Feedback (FB) Onset (0ms) as well as 500ms and 1,000ms after. These time points were also chosen based on previous TMS work using similar timings to show reward- and punishment-induced excitability changes during feedback processing^41,42^.

### EMG system and neuronavigated TMS

Electromyography (EMG) data from the right FDI muscle belly were recorded through a single bipolar electrode connected to a 2-channel Delsys Bagnoli (Delsys ®) system, itself connected to a Micro 1401 data acquisition unit (Cambridge Electronic Design). The EMG data were acquired with Signal (Cambridge Electronic Design, v6.05) at a sampling rate of 10,000Hz for epochs of 500ms (200ms pre-trigger time). The EMG data were high-pass and low-pass filtered at 20Hz and 450Hz, respectively, with a notch at 50Hz. The reference EMG electrode was positioned on the proximal olecranon process of the right ulnar bone. The EMG data recorded were exported in MatLab, which were analysed using an automated custom-built script.

Neuronavigated TMS pulses were delivered through a single figure-of-eight 70mm Alpha Flat Coil (uncoated) connected to a paired-pulse BiStim^2^ stimulator (MagStim, Whitland, UK). BrainSight (Rogue Research; Montreal, Canada) was used to ensure reliable coil positioning during every experiment and session^81^. The coil was positioned at a 45° angle in a posterior-anterior axis over the FDI motor hotspot of the left M1, defined as the area where MEPs of maximal amplitude could be reliably elicited with suprathreshold pulses in the right FDI. The resting motor threshold (RMT) was defined as the % of maximum stimulator output to induce 5 out of 10 MEPs of at least 50µV of peak-to-peak amplitude^82^. For every participant, the FDI motor hotspot, the RMT, and the test stimulus intensity (see below) were assessed at the start of every experiment and session. For both sessions of the ICF and SICI experiments, the average RMTs (± SEM) were 49 ± 2% and 51 ± 2% of the maximum stimulator output, respectively. Bivariate correlations revealed that RMTs between the Movement Preparation and Feedback Processing sessions were highly correlated in the ICF (R = 0.933, *p* < 0.0001) and SICI Experiments (R = 0.973, *p* < 0.0001), suggesting that RMT assessment for a given participant was stable across sessions.

The used TMS parameters are shown in Figures 1C and 1D as well as 3C and 3D and were the same for both the Movement Preparation and Feedback Processing sessions. ICF and SICI were *respectively* induced by delivering conditioning pulses at 90% and 70% of the RMT^83^. Inter-pulse intervals of 15ms and 3ms were selected to induce ICF and SICI^36^, respectively. For both ICF and SICI, test stimuli (TS) intensity was calibrated to induce motor-evoked potentials (MEPs) of ± 1mV at rest. For both sessions of the ICF and SICI experiments, the average TS intensities (± SEM) were 58 ± 2% and 62 ± 3%, respectively. Bivariate correlations revealed that TS intensities between the Movement Preparation and Feedback Processing sessions were highly correlated in the ICF (R = 0.927, *p* < 0.0001) and SICI Experiments (R = 0.945, *p* < 0.0001), suggesting that assessment of TS intensity for a given participant was stable across sessions.

### Trial rejection

Behavioural trials where RTs were below 200ms were rejected from the analyses, as these trials were indicative of premature responses. To prevent the contamination of muscle pre-activation in TMS data, TMS trials where the average root mean square of the FDI EMG amplitude exceeded 100µV in the 50ms before the TMS stimulator was triggered were removed from analyses (similar to ref ^84^). Finally, for the Movement Preparation session only, the TMS pulses at 400ms that were delivered to a latency > 100% of RT were rejected, since TMS pulses would then be delivered during execution (outside of preparation). Across both Experiments and sessions, this resulted in a total rejection of 2.7% of all trials (936 out of 34,560 trials).

### Number of TMS trials per condition

For both the ICF and SICI experiments, a total of 18 TMS trials were delivered per TMS variable (CSE, ICF or SICI), per level of Incentives (Reward, Neutral, Punishment), and per Time Point (GoCue, Early RT, Late RT or FB Onset, 500ms, 1,000ms). This resulted in a total of 324 TMS trials (out of 432 behavioural trials). For the Movement Preparation session only, the TMS data were first pooled according to the percentage of the RT period at which pulses were delivered (Figure 1A). This was because reaction times (RTs) were variable, but TMS was delivered at fixed time points, implying that TMS pulses were unlikely to be systematically delivered at the same percentage of the RT period across trials. TMS pulses delivered at a latency below and above 50% of the RT were pooled together and defined as Early RT and Late RT, respectively. Pulses delivered at GoCue onset were not pooled differently, since their delivery precisely coincided with the start of the RT period. For the Feedback Processing session, TMS data were independent of RT and were thus pooled according to their delivery time point. Overall, the number of trials reported below exceed current guidelines to reliably assess excitability changes (TMS without neuronavigation^85^: 30 trials for ICF and 26 for SICI; TMS with neuronavigation^86^: 25 trials for ICF and 20 trials for SICI), confirming that the effects of Time Points and Incentives of both Experiments are supported by reliable MEP measurements.

For each session of each Experiment, there was as many CSE trials as there were ICF and SICI trials included in the final analyses, suggesting that the CSE measurements were reliable and offered a stable baseline for normalisation^85,86^. Concerning the evaluation of ICF during Movement Preparation, per participant, there was an average of 53 ± 0.6, 55 ± 1.9, and 38 ± 2.0 CSE trials for the GoCue, Early RT, and Late RT time points (collapsed across incentives), respectively. In that same session, per participant, there was 52 ± 0.6, 52 ± 0.5, and 52 ± 0.7 CSE trials for the Reward, Neutral, and Punishment conditions (collapsed across time points), respectively. Concerning the evaluation of SICI during Movement Preparation, per participant, there was an average of 52 ± 0.9, 55 ± 2.8, and 36 ± 2.1 CSE trials for the GoCue, Early RT, and Late RT time points (collapsed across incentives), respectively. In that same session, there was an average 51 ± 0.9, 52 ± 1.0, and 51 ± 0.9 CSE trials for the Reward, Neutral, and Punishment conditions (collapsed across time points), respectively.

Concerning the evaluation of ICF during Feedback Processing, per participant, there was an average of 52 ± 0.8, 52 ± 0.7, and 52 ± 0.7 CSE trials for the FB Onset, 500ms, and 1,000ms time points (collapsed across incentives), respectively. In that same session, per participant, there was 53 ± 0.7, 52 ± 0.7, and 52 ± CSE trials for the Reward, Neutral, and Punishment conditions (collapsed across time points), respectively. Concerning the evaluation of SICI during Feedback Processing, per participant, there was an average of 53 ± 0.4, 54 ± 0.4, and 53 ± 0.2 CSE trials for the FB Onset, 500ms, and 1,000ms time points (collapsed across incentives), respectively. In that same session, per participant, there was 53 ± 0.4, 53 ± 0.4, and 54 ± 0.3 CSE trials for the Reward, Neutral, and Punishment conditions (collapsed across time points), respectively.

Concerning the evaluation of ICF during Movement Preparation, per participant, there was an average of 53 ± 0.6, 61 ± 2.9, and 35 ± 2.3 ICF trials for the GoCue, Early RT, and Late RT time points (collapsed across incentives), respectively. In that same session, per participant, there was 50 ± 0.8, 50 ± 0.7, and 50 ± ICF trials for the Reward, Neutral, and Punishment conditions (collapsed across time points), respectively. Concerning the evaluation of SICI during Movement Preparation, per participant, there was an average of 47 ± 2.1, 47 ± 2.8, and 38 ± 2.0 SICI trials for the GoCue, Early RT, and Late RT time points (collapsed across incentives), respectively. In that same session, there was an average 44 ± 1.6, 44 ± 2.0, and 44 ± 2.1 SICI trials for the Reward, Neutral, and Punishment conditions (collapsed across time points), respectively.

Concerning the evaluation of ICF during Feedback Processing, per participant, there was an average of 52 ± 0.8, 52 ± 0.7, and 53 ± 0.6 ICF trials for the FB Onset, 500ms, and 1,000ms time points (collapsed across incentives), respectively. In that same session, per participant, there was 53 ± 0.7, 53 ± 0.6, and 52 ± 0.8 ICF trials for the Reward, Neutral, and Punishment conditions (collapsed across time points), respectively. Concerning the evaluation of SICI during Feedback Processing, per participant, there was an average of 52 ± 1.1, 51 ± 1.2, and 50 ± 1.4 SICI trials for the FB Onset, 500ms, and 1,000ms time points (collapsed across incentives), respectively. In that same session, per participant, there was 51 ± 1.1, 51 ± 1.2, and 51 ± 1.3 SICI trials for the Reward, Neutral, and Punishment conditions (collapsed across time points), respectively.

### Dependent variables

The behavioural dependent variables were as follows. Reaction time (RT) was defined as the time difference in milliseconds between GoCue onset and the first finger press (keypress). Movement time (MT) was defined as the time difference in milliseconds between the first and last finger press (keypress). Accuracy (success rates) was defined as a binary variable denoting if participants executed the correct sequence within the allowed execution time (Figures 1A and 3A). The TMS dependent variables were calculated as MEP peak-to-peak amplitude upon delivery of single (CSE) and paired TMS pulses (ICF or SICI). For single pulses (CSE), MEP peak-to-peak amplitudes were calculated as (non-normalized) values (in mV) and averaged separately for each level of time point, incentive, session, and experiment. For paired pulses, the MEP peak-to-peak amplitudes measured on ICF and SICI trials were calculated for each level of time point, incentive, session, and experiment. Then, these individual trials were separately normalised as a percentage (%) of their corresponding average CSE. Subsequently, the normalised ICF and SICI trials were averaged separately for each level of time point, incentive, session, and experiment.

### Statistical analyses

The main analyses were conducted using generalised mixed linear models^87,88^, with a gamma distribution to account for the positive continuous skewness of the RT, MT, and MEP data^39^. For the Accuracy (success rates) variable, generalised mixed linear models with a binomial distribution were conducted to deal with their binary nature (on a trial-per-trial basis). For each dependent variable of both the Movement Preparation and Feedback Processing sessions of both the ICF and SICI Experiments, the maximally complex random effect structure (random intercepts for Participants and random slopes for all of the Fixed Effects and Interactions, wherever the data allowed their inclusion) that minimised the Akaike Information Criterion (AIC) was chosen to analyse the results^89^. For the Movement Preparation session, the fixed effects were Incentives (Reward, Neutral, Punishment), Time Points (GoCue, Early RT, Late RT) and Initiating Finger (Index, Little). For the Feedback Processing session, the fixed effects were Incentives (Reward, Neutral, Punishment) and Time Points (GoCue, Early RT, Late RT). The fixed effects were the same for both the ICF and SICI Experiments. P values below 0.05 were determined as statistically significant. The Benjamini-Hochberg (1995)^90^ correction was used to correct p values for multiple comparisons. To report statistics, the mean ± 1 SEM was used throughout. All analyses were conducted in JAMOVI (version 2.3.16)^91^.

## Acknowledgements

This work was funded by the *Fonds de Recherche du Québec – Nature et Technologie* (Québec, Canada), European Research Council Starting Grant (MotMotLearn 637488) and Proof of Concept Grant (ImpHandRehab 872082).

## Data availability

This work’s data are freely available and can be obtained as an Excel spreadsheet from the following URL: https://osf.io/96u7q/

